# A novel phage satellite class induces prophage excision in *Mycolicibacterium aichiense*

**DOI:** 10.64898/2025.12.02.691453

**Authors:** Heather L. Qian, Anne N. Roman, Sudip Paudel, Grace E. Hussey, Kate B. R. Carline, Margaret S. Saha

## Abstract

Phage satellites are mobile genetic elements that parasitize a helper phage to complete their life cycle. While phage satellites are widespread in diverse bacterial hosts, none have been isolated in *Mycobacteriaceae*. Here, we report the first phage satellites isolated and characterized in *Mycobacteriaceae*, Extracellular Prophage-Inducing Particles (EPIPs). EPIPs induce a helper phage—HerbertWM, a *Mycolicibacterium aichiense* prophage—upon infection. Genomic sequencing of thirteen isolates revealed a notably small genome size and high sequence similarity between EPIPs, suggesting they are a distinct class of phage satellite. EPIPs were categorized into groups based on Virus Intergenomic Distance Calculator similarity scores, gene content, and synteny. Gene content is largely conserved across the isolates, though EPIPs notably lack capsid proteins, tail proteins, and holins, suggesting EPIPs hijack the machinery of HerbertWM to replicate, assemble, and lyse the host. Furthermore, the unique gene content of EPIPs compared to other phage satellites, particularly the presence of a tape measure protein and absence of antirepressors, suggests they may exhibit unique mechanisms of assembly and prophage induction. Transmission electron microscopy of EPIPs reveals that they not only feature smaller capsids compared to HerbertWM, but that they feature longer tails of variable length, further suggesting a unique mechanism of tail hijacking that the tape measure protein may facilitate. Additionally, greater EPIP protein similarity to prophage proteins as opposed to free phage proteins suggests a functional or evolutionary relationship may exist. The synteny among EPIPs and the unique gene content suggest EPIPs are a distinct class of phage satellites that induce *Mycobacteriaceae* prophages.

## Main Text

Phage satellites (satellites) are mobile genetic elements (MGEs) that hijack helper phage machinery to complete their life cycle. They can exist extrachromosomally as a plasmid or phagemid, integrated in the bacterial host genome, or as a free infective particle [reviewed in 1, 2]. These interactions create a parasitism-mutualism continuum between the helper phage and satellite; satellites benefit the helper phage by blocking non-helper phage infection and competing MGE transfer while simultaneously inhibiting helper phage replication. Satellites also confer selective advantages to the host: immunity against non-helper phages, increased virulence, and transduction of bacterial genes [1, 2].

Three classes of satellites—P2-P4-like systems, phage-inducible chromosomal islands (PICIs), and PICI-like elements (PLEs)—have been well characterized. All remodel helper capsids and have DNA sequences recognized by a native or hijacked helper phage terminase [1, 2]. However, satellite classes display differences: mutual induction (satellite and helper phage inducing the other) in P2-P4-like systems, independent replication in PICIs, and complete inhibition of helper phage replication and acceleration of host lysis in PLEs [1,2]. Furthermore, satellites are highly diverse with discoveries of novel classes growing rapidly. For example, capsid-forming PICIs (cf-PICIs), MiniFlayer, MulchRoom, and Phanie encode capsid proteins; phage-inducible chromosomal minimalist islands (PICMIs) and virion-encapsidated integrative mobile elements (VEIMEs) package concatemeric genomes; MiniFlayer and Phanie attach to their helper phages for simultaneous infection; and ϕAH14b and its helper lack tails. Additionally, tycheposons encode phylogenetically distinct tyrosine recombinases [1, 3, 4]. Despite this diversity, satellites all share small genomes and rely on helper phage machinery [1, 2].

Here we report yet another novel class of satellites named “Extracellular Prophage-Inducing Particles” (EPIPs) due to their ability to induce the excision of the HerbertWM helper phage, a *Mycolicibacterium aichiense* prophage, making EPIPs the first satellites experimentally isolated from *Mycobacteriaceae*. EPIPs were isolated from soil near Jamestown, Virginia over a two-year period using standard procedures for phage isolation, and thirteen isolates were selected for DNA sequencing and analysis (Supplementary Methods).

Assembled reads for all samples revealed two circular contigs (Supplementary Methods). One, identical across all samples, was identified as the prophage HerbertWM. A second 11-kb contig, the EPIP, varied among samples, showing sequence similarities ranging from 58.1% to 99.8% determined using Virus Intergenomic Distance Calculator (VIRIDIC) [5] (Fig. 1D). Stoichiometric ratios between EPIP and HerbertWM reads varied between samples (EPIP-to-HerbertWM ratio: lowest 1:1, highest 725:1).

**Fig 1.**
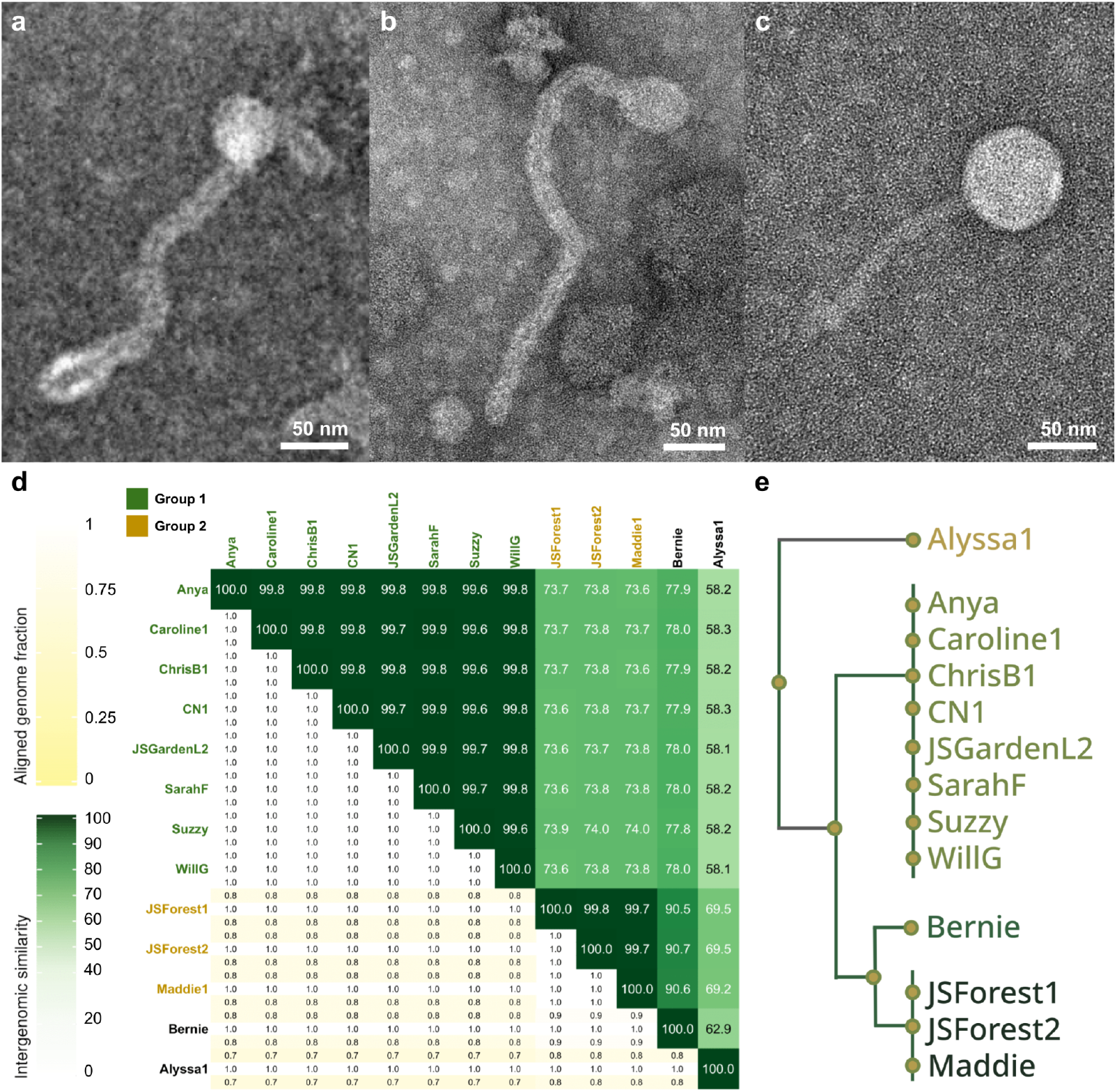
EPIP morphology and genomic similarity. **a** Alyssa1 (mean capsid diameter 51.2 ± 1.2 nm, n = 3; mean tail length 384.5 ± 119.4 nm, n = 3). **b** Bernie (mean capsid diameter 32.0 ± 4.1 nm, n = 4; mean tail length 283.8 ± 42.4 nm, n = 3). One particle was excluded from tail length measurements due to likely tail breakage. **c** The HerbertWM helper phage, a *Mycolicibacterium aichiense* prophage. **d** Genome similarity generated by VIRIDIC (mean percent within groups 99.76 ± 0.01, Group 1 vs Group 2 73.75 ± 0.2, Alyssa1 vs all other EPIPs 58.20 ± 1.4, Bernie vs all other EPIPs 79.85 ± 2.2). **e** Phylogenetic tree of EPIPs generated by Clustal Omega Multiple Sequence Alignment. Each branch endpoint represents a single EPIP, and each branch represents a group.

EPIPs were categorized into two groups defined by a VIRIDIC similarity score greater than 95%—the VIRIDIC species threshold [5], identical protein function, and synteny, with each group comprising a distinct branch on the phylogenetic tree generated with Clustal Omega [6] (Fig. 1D-E). Within each group, EPIPs share ∼99.8% VIRIDIC similarity, while between groups, EPIPs share ∼73.7% similarity. The uncategorized EPIPs, Alyssa1 and Bernie, respectively share ∼58.2% and ∼79.9% similarity with all other EPIPs and were included in comparative analyses (Fig. 1D). Additionally, all EPIPs lack homology with HerbertWM (∼8.5% nucleotide coverage, ∼73.1% identity) determined using MegaBLAST [7].

Genome annotations were determined using multiple metrics including DNA Master v5.23.6 [8, Glimmer v3.02 [9], GeneMark v2.5 [10], BLAST [7], and HHPred [11]. BLASTp [7] was used to align each EPIP with proteins in NCBI and PhagesDB, an actinobacteriophage database [12]. Foldseek [13] and DALI [14] were used to align EPIP protein structures predicted by AlphaFold 3 [15] with proteins in Protein Data Bank [16] (Supplementary Methods). All EPIPs encode a terminase small subunit, head-to-tail adaptor, head-to-tail stopper, serine integrase, helix-turn-helix DNA binding domain, scaffolding protein, major capsid protein, tape measure protein, and four conserved hypothetical proteins. However, all EPIPs lack genes critical for phage function such as capsid proteins, tail proteins, and holins, suggesting EPIPs parasitize HerbertWM to replicate, assemble, and lyse the host (Fig. 2A).

**Figure 2.**
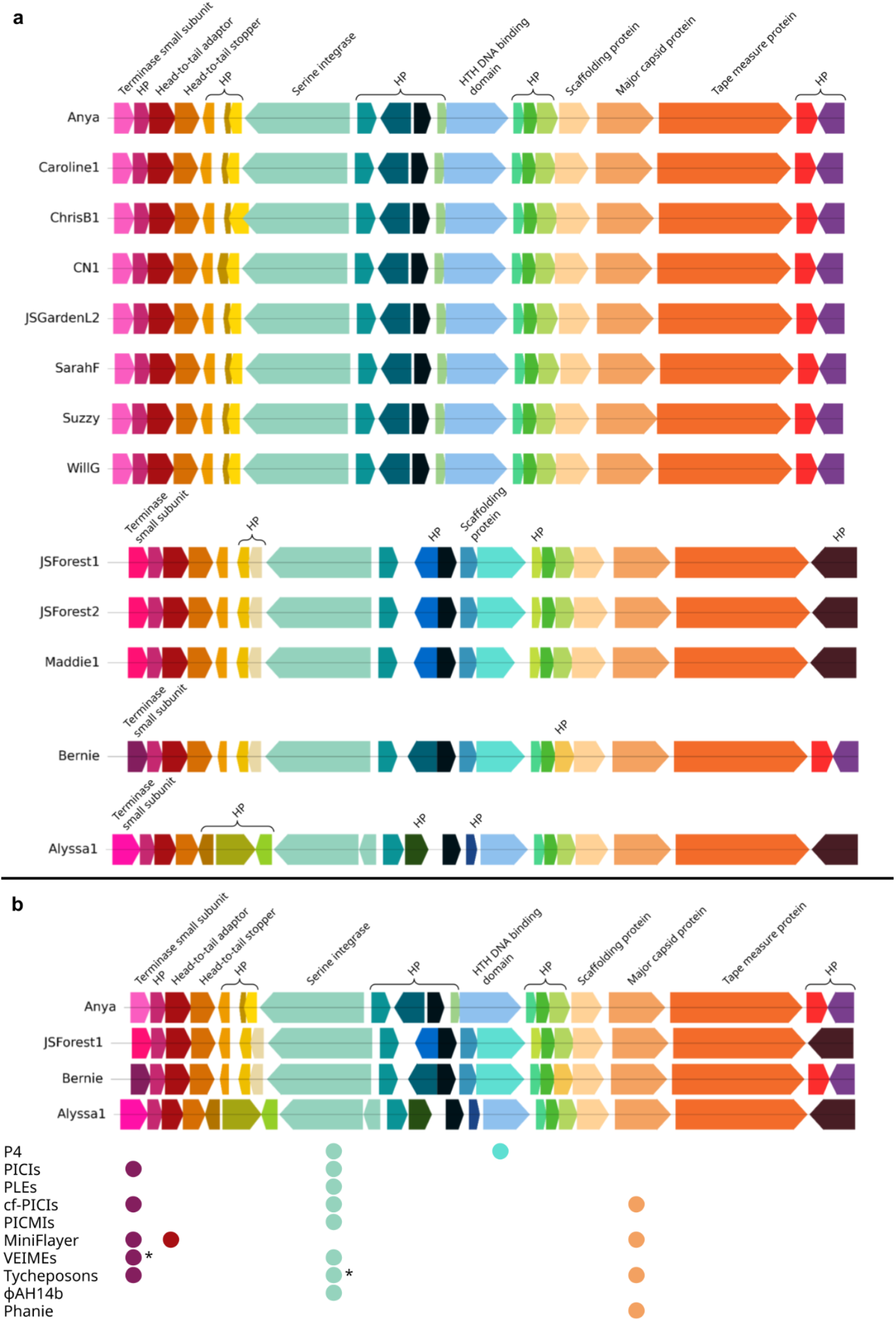
EPIP gene functions. **a** Alignment of EPIP proteins, with proteins of the same color being homologous (≥ 70 % amino acid coverage, ≥ 30 % identity; Supplementary Methods). **b** EPIP proteins common to other phage satellite classes. The column of each dot represents the EPIP protein, and the row represents the phage satellite class that contains a similar protein. * 8% of VEIMEs encode terminases, and tycheposons typically encode tyrosine integrases.

The presence of a tape measure protein (TMP) is notable, as TMPs are not present in satellites with only a recent preprint proposing that PLE11 encodes a TMP-like protein [17]. The TMP may impact EPIP morphology, as transmission electron microscopy of two EPIP isolates, Alyssa1 and Bernie (Supplementary Methods), revealed they feature tails that are longer and more variable in length (∼385 and ∼284 nm respectively) than HerbertWM (∼134 nm). This suggests the TMP may facilitate a unique mechanism of tail hijacking. Additionally, Alyssa1 and Bernie possess smaller capsid diameters (∼51 and ∼32 nm respectively) than HerbertWM (∼67 nm), whose capsid diameter and tail length are common to mycobacteriophages (40-80 nm and 135-350 nm respectively) [18] (Fig. 1A-C).

Despite significant similarities, EPIPs exhibit variation across groups; a scaffolding protein and many hypothetical proteins are not conserved across all groups (Fig. 2A).Additionally, although intergenic regions are well-conserved within groups (100% median VIRIDIC similarity), intergenic regions across groups are significantly less conserved (86.1% median VIRIDIC similarity, p = 2.23 × 10^-68^). Interestingly, regions between genes transcribed in opposite directions are more conserved across groups than regions between genes transcribed in the same direction (p < 0.05), suggesting conserved regulatory architecture. Notably, alignment of EPIP genes against proteins in NCBI and PhagesDB demonstrated higher sequence similarity to prophage proteins (p < 0.001) as confirmed using PHASTEST [19] and DEPhT [20], suggesting a close evolutionary or functional relationship between EPIPs and prophages (Supplementary Methods).

The striking similarities between EPIPs across groups—synteny, high VIRIDIC similarity, and homology to prophages—despite EPIPs being isolated from a wide geographical and temporal range suggests EPIPs are a new class of satellites that induce *Mycobacteriaceae* prophages. The similarity of EPIP proteins with prophage proteins and the ability of EPIPs to induce prophage HerbertWM suggests that EPIPs share a functional or evolutionary relationship with prophages. While EPIPs share commonalities with other satellites, such as small genomes and a lack of proteins required for an independent life cycle [1, 2], EPIPs display distinct characteristics as well. For example, EPIPs induce the helper rather than vice versa. Moreover, the distinct gene content of EPIPs, particularly their encoding of a TMP and lack of canonical antirepressors, may suggest unique mechanisms of assembly and prophage excision (Fig. 2B). Taken together, the data presented here suggests EPIPs are a distinct class of *Mycobacteriaceae* satellites whose novelty invites further investigation of their mechanisms of action.

## Acknowledgements

We thank the many high school students from Jamestown High School who participated in the discovery of these satellite phages. We also thank Taiana James, Mitchell K. Doherty, Mahima Shijo, and Rebecca D. Zheleznyak and the many others for assistance with various aspects of EPIP discovery and characterization. We thank the Pittsburgh Bacteriophage Institute for sequencing and assembling HerbertWM and the Bernie EPIP isolate.

## Data availability

Genome sequences used for analyses were previously deposited in NCBI GenBank under the following accession numbers: MN224566, OR387110, OR387111, OR387112, OR387113, OR387114, OR387115, OR387116, OR387117, OR387118, OR387119, OR387120, OR387121. The genome sequence of the Bernie EPIP has been submitted to GenBank and is awaiting an accession number; the sequence is available in Supplementary Information.

