## Supplementary Information for "A novel phage satellite class induces prophage excision in *Mycolicibacterium aichiense*"

### Supplemental Methods

All statistical tests were performed using SciPy v1.15.3 [1] in Python v3.10.

#### EPIP isolation

As previously described [2], after inoculation of 5 mL of soil with 25 mL Middlebrook 7H9 media, 3 mL AD supplement, and 1 mL liquid *M. aichiense* NCTC 10820 (ATCC 27280) culture, samples were incubated at 37ºC with shaking (250 rpm) for three days. Following centrifugation at 3,400 rpm for 15 minutes, the culture was filtered through a 0.22 um PES filter. The resulting lysate was incubated with *M. aichiense* for 20 minutes at room temperature, then the infected culture was plated using Middlebrook 2X 7H9 top agar on LB plates (15 g/L agar, tryptone 10 g/L, yeast extract 5 g/L, NaCl 10 g/L) and incubated for two days. Resulting plaques were purified through two additional rounds of plating before lysates were collected by flooding webbed-lysis plates with 5 mL phage buffer (1 mM $CaCl_2$, 10mM Tris, 10 mM MgSO4, 0.4% w/v NaCl) and incubating the flooded plates for 14 hours at 20ºC. Collected lysates were filtered using a 0.22 um PES filter.

#### TEM

Lysate was either collected by flooding a webbed-lysis plate (Bernie) or by extracting spots resulting from a spot assay and placing the plug in 500 uL phage buffer at 4ºC overnight (Alyssa1). Following a standard protocol for TEM grid preparation [2], 4 mL lysate was filtered using a 0.22 um PES filter into two 2 mL tubes. Following centrifugation at 13.3k rpm at 4ºC for 1 hour, supernatant was decanted and pellets were resuspended in 50 uL phage buffer. After incubating at 4ºC for 1 hour, the two 50 uL volumes were combined. TEM grids were prepared by placing 10 uL lysate onto carbon-formvar-coated copper grids, and EPIPs were allowed to attach for 7 minutes. The grid was rinsed with 10 uL nuclease free water twice before being stained with 10 uL 1% uranyl acetate for 2 minutes.

#### DNA extraction

To increase the titer of the lysate, polyethylene glycol (PEG) precipitation was performed [3]. 1% chloroform was added to an infected culture of *M. aichiense* cells and then incubated for 90 minutes at 37°C. A sample of this culture was then centrifuged at 3500 rpm for 5 minutes and was followed by filtration with a 0.22 µm PES filter. To 95 mL filtered lysate, NaCl was added to 1M followed by 10% volume PEG 8000, and the sample was incubated overnight at 4°C with stirring. The sample was then centrifuged at 4°C for 10 minutes at 5500 x g and supernatant was removed. The remaining pellet was resuspended in 5 mL phage buffer.

To extract DNA [4], 12.5 uL 1M $MgCl_2$, 14 uL nuclease mix (4 uL 200 U/mL DNAseI and 10 uL 10 mg/mL RNase A was added to the PEG-treated lysate. The tube was vortexed to mix then incubated at room temperature for 30 minutes. Then, 40 uL 0.5 M EDTA, 5 uL 10 mg/mL Proteinase K, and 50 uL 10% SDS were added, and the tube was vortexed before incubating at 55ºC for 1 hour, vortexing twice during this period at 20 minute intervals. Samples were divided into 500 uL volumes and DNA was extracted using 500 uL PCI (25 phenol : 24 chloroform : 1 isoamyl alcohol). The tube was inverted to mix, then centrifuged at 13,000 rpm for 5 minutes. After transferring the upper phase to a new tube, 500 uL CI (24 chloroform : 1 isoamyl alcohol) was added, and the tube was centrifuged at 13,000 rpm for 5 minutes after inverting to mix. The upper phase was transferred to a new tube, and 50 uL 3M sodium acetate and 1 mL ice cold 100% ethanol were added. Following incubation on ice for 7 minutes and centrifugation at 13,000 rpm for 10 minutes, supernatant was decanted and the pellet was washed with 500 uL ice cold 70% ethanol. Samples were centrifuged again at 13,000 rpm for 10 minutes, and supernatant was discarded. Pellets were air dried for 20 minutes before being resuspended in 50 uL nuclease free water. DNA was quantified using a Nanodrop spectrophotometer and visualized using agarose gel electrophoresis.

**Genomic sequencing**

EPIP genomes were sequenced using an Illumina MiSeq sequencer and reads were assembled using Newbler v2.9 (RRID:SCR_011916). The final assembly was determined using Consed v29.0 [5] at the Pittsburgh Bacteriophage Institute or using CLC Genomics Workbench v22.0 (QIAGEN). The starts of EPIPs' circular genomes were aligned using CLC Genomics Workbench v22.0 (QIAGEN).

**Protein sequence alignment analysis**

BLASTp [6] was used to align each EPIP protein with proteins in NCBI's ClusteredNR database and PhagesDB [7]. Many EPIP proteins aligned to a prophage protein in ClusteredNR rather than a free phage protein in PhagesDB, so the bacterial species was analyzed using PHASTEST [8] and DEPhT [9]. Using BLASTp, EPIP proteins were then aligned against prophage proteins identified by PHASTEST. Four EPIPs were selected for further analysis: WillG, JSForest2, Bernie, and Alyssa1. Bit scores were recorded for all alignments against proteins in NCBI's ClusteredNR, PhagesDB, and PHASTEST-identified prophages. Because 44.6% of EPIP proteins did not align to bacteria containing prophages, this set of bit scores was excluded from further analysis. Using a Wilcoxon signed-rank test, EPIPs were determined to exhibit significantly higher homology to proteins in ClusteredNR than in PhagesDB ($p < 0.001$).

**Protein structural alignment analysis**

One EPIP from each group and the two uncategorized EPIPs were selected for analysis. EPIP protein structure was predicted using AlphaFold 3 [10], and predicted structures were aligned with structures in Protein Data Bank [11] using Foldseek [12] and DALI [13]. Alignment quality was evaluated using overall fold similarity, percent coverage, and the context of the EPIP protein within the EPIP genome. For Foldseek results, overall fold similarity was determined using the overlay function in ChimeraX v1.10.1 [14], while for DALI results, overall fold similarity was determined using side-by-side comparison.

**Intergenic regions analysis**

Intergenic regions were defined as any region outside of an annotated gene, with sets defined as intergenic regions between proteins of the same function across EPIPs. Each set of intergenic regions was manually extracted from genome sequences based on start and stop positions of genes. Within each set, pairwise similarity of intergenic regions was evaluated using a Virus Intergenomic Distance Calculator similarity score [15]. Then, to determine whether similarity scores were higher for intergenic regions within groups than between groups, pairwise comparisons of similarity scores were performed using Mann-Whitney U tests. Then, a global Fisher p-value of $2.23 \times 10^{-68}$ was calculated with a similarity threshold of 95 to combine all Mann-Whitney U scores.

To investigate the conservation of intergenic regions across groups, intergenic regions were then marked as positioned between genes transcribed in opposite directions or in the same direction. A subset of similarity scores was created by selecting similarity scores of intergenic regions from different groups. Then within this subset, a Mann-Whitney U test was used to compare the median similarity of regions between genes transcribed in opposite directions (95.87%) versus regions between genes transcribed in the same direction (84.16%) (Mann-Whitney U = 13134.50, $p = 1.82 \times 10^{-2}$).

**Generation of Figure 2A**

Genome maps were generated using gene start and stop sites from exported annotations. A previously described homology threshold of ≥ 30% amino acid identity [16] and ≥ 70% amino acid coverage was used to minimize alignments solely due to shared domains [17].

**Bernie Genome Sequence**
TGCGGGCGGGGCATCGGGTGCGACTTAAAGCCCTGTTTTTCGAGAGTCAGTAAGTTCCGGTCAGCGAAC
GAGAGCTGACCTAAGTTCAACTTAGGCCAGGTCCAATCGTCGCAGATCAGAACAAGAGTGTGTGCAATG
TACCCACCCCTGCCAGCAAATTACATCGTGAATCACGATGGTATTCCACGCTCATCAGCACAACCCCAG
GTCAACCCTGAGGTGATCACTAATCACCCCTTCGTGACCTGCAAAAACGTCCATTCGTCGGTTGAACCG
ATGTATGCCCCAGGAAGGAACCCCAGATTGAGAGCGCCAACCGGCCTACAAGCCAGCGGAAAGCGGCTC
TGGAAGTCGATTACCTCAGAGTTCGACCTGTCGGCTGATCCCGACAAGGCCGAGTTGCTGTTCCAGGCA
TGCCGAACCGCTGACCAGATCGCGGAGCTGGACGAGGCAGCCGCCGAGGCACCGCTGACCGTCAAGGGC
TCGATGGGCCAGCCGGTGATCTCGCCGTTCATCGCAGAGGCCCGCGTCCAACGCGGTCTCCTGGCTCAG
CTCTTGTCGCGGATGAACTTCGAGGATGCGATCGATGAGTAAGGCCGGTCGGAAGACCAAGCGGCATCG
CAACGCATCCGCTCCCGCCGTAGACGCGGGAGCTGTTTTCCTGCGGCTGATGAAGCAGCGGTTGACCAA
CCCCGACCCGCGCTCGCACGCGGAGTTCCTCGCAGACATGCGAGCCAAGCAAGCGGCAGGACGGCTGGC
CAACCCGGCCACTGCCGCACTACTCGACCGACTAGAGAGGGCCATCAAATGACCTACGCAACAGCCGCT
GACATAGCCGCGCTCCTGGCGCGGGAGCTGGACACCGCCGAGACGGCGCTGGCCGAGCGCCGACTGGCT
CAGGTAGAGCGAATGATCCTGCGCCGCATACCTGATCTTGCCGACCAGATCGCCGCTGGCGACCTGGCC
GAGGCCGACGTGATCGACGTCGAGGCCGAGGCCGTCTACCGCGTCATGCGGAACCCTGACGGGCTCTAC
AGCGAGCAGGACGGCCAATACGGATACCAGCTCTCCCGCGAGGCTGCCGACAACAGCCTCCGCGTCACC
ACCGAGGAGTGGCAGACCCTCGGCATCAAGCCGAGCAAGCTGTTCTCGATCGCTCCGCGTATCGGAGGC
TTCGCGTGAGCCTTCTACGACGTGGAACTGAGACCGTCACGGTCTACCCCGAGGTGACCGTGACCGACA
GCGACGGCAACAAGTTCACTCGCCCCGGTTCGGTCGGAATCGTCGTCAAAGCGTCGGTTCAGCCCCTAG
GTCTGCTCGGCGCTCCCGCCGAGAACACGGACGGGGGATTCAACACCCAGAGCCGATACCGGCTGCGGC
TGGCAGGCGGATGGCCAACCGGCGGCGGCATTCTGGGGGCTCAGGCACGGGTCGAATGGCGCGGCAAGC
GGTACTCGATCGAAGGCGATGCCCAGTTGCACACCGGCAGCCCCAGGACCGCCCACGCGGTCTACACGA
TGGTTCGGAGCTGACAGCGCCGACTCGCTGATGAAGTTCAGCGACTGGTTGGGGCCACTACCCTGATTC
AGGGTAGTGCGCTGATCTGCGTCGATGAGGGCTCAGTCGTCCGGAATGCCGTCCGTAGTTCTTGCAGGC
CACCGGGGGGTGTCCACAGTGTGTACACCCTTCTCGTCTTCGTCCAGAAGAGCGAACGACGACCGGCTC
ACCCCCAACGCTTGCAGCACGTCAGCAGCCGCTCCTGCGGGCGGCTCTTCCCGGCCTCACCGTCGCCTC
CTCAGACCCCGGCCCCGAGCACCGTCAGTTCCGAGAACGGGCCCGAGGCTGTGCGGTACCACGTCCAAG
AGTCGCAGGTCAGAAGGGGTGAGCGAATCCCTCACCCCCAGTCGACCACGGCTGGCCGTCCGGTCGGCG
TAGCCACGTTCGACCGCATGATCAGCCGGTAGAGGCTGCTCTCACTGACCAGGATCTGGTTCCGGTACT
GCCTGCCCTCGTGAATCGCGAGGTCAGCTCGCTGATCAGGGAAAACGTGCTGGTTCACCGCCAGGGGTA
CCGATTCGGTACTGCTGGGCCTGACCTGCGACTCCTGAACGGGGTGAACGATTCGTTATGACCCGCGCT
GATGATGGAAACCTGCTGGTCACCACCGGGGGTGCCGATTAGGCACCTCCCTTTCTCGTCATCATCGAG
GGACGAGATGGCCATGCTCACCGGGCGGGGGGTCATCAACTGGATGACCCCTTGCACGTCCTTGGCGAC
GAACCACGGCTCGCCGTCGACAGGAACGACGCGGCGGCTGCTCAGTCCTCCGGGATGCCGAGCTTGCGC
CGCTCGGCGGCGGCCCCCGCATGCCACCGCTCGATCTCCTCGGGAGAGCTGTTCTCGTCTACCTGAGCC
CAATCGGGCAGCGGCAGCCGGTCCCCGAGCCTCAGCGGATGAAGCAGTAACTGGCCGTCGCTGACCCCG
GCACGAACCTTGAACCCGATGCGCTTCAACAGAGCCCGACGTTCATCGGGGTTGTCGGCGGTTCTCTCC
AGCTCCGAACGGTATGTCTTGCCGGTCGGTCGGATCTCGTACCGACCCTCCTGGGAGGGCAACAACTCC
AGCTCTGCGATTCGAGCGTCCAACGCGCTGAGTTGGCGCTGGAGAATGTCTTTCGCCGTTCGGGAGGTC
GCGGTACCAGCCGTAGCCGAAAGCTCGGTGTACGCCGCTACAGCCTCTCGCAGAGCGGCCTCGTTCGAG
TCGCCGGGCACCCAGACCCGCTCGGTCACCGGTTCGTCTGCGAGAGCGCTCATCAGCACGCGGTCAACC
TCCTCCTCAACGGTCTCCGCTCGGACCGAGGAGCAATTTCCGCTGTAGCACCGGTAATAGCGATACCGC
TTGCCGTTCTGTGTTCTGGCAGAAGAGGTCAGCGGTCCCTCGCAGAACCAGCACTCGATGAGACCCGAC
AACGGCGCGGTCTCCAACCGCTGATAGGAACCCTTGGCGACGCGGGCCAGCTCGGCTCGGATCAGCTCT

CGCTCGTCGTCTGTGACGAGAGGCTCTCCGAGCTGTATCGGCATACCCTGCTCGTCGCGCACGGTCTGC
CCGCCCAAGTGGGCGTGGCCGACCAGTGCGGGTGACCTCAGCATGTCGTGCAACGGGGCGCGCTGCCAT
TTGCCGCCCCTGGCAGGCTTTCCCATCGACAGGCGGTAGTGATCAGCCGGTGACGGGATGCCGTCCTTG
TTCAGCCGGTCGGCGATCTGAGCGAGCGAGTTCTGTTGCAGCAGTTCCTCTACGATCCGGCGCACCACC
TTGTGCGCCACTGGATCGATCACCAGAATCTTGCCCGACCCGGTCGGGCTGTTCGCGGTCTTGTAGCCG
TACGGCGGCTTGCCTCCCGGCCAGCGGGCGGTCTCGCGCAGCTTGCGCCGGGAGCTGATGGTTCGCTCC
CTGATCGCCTCCAGCTCGCCCTCGGCCAGGAACCCGATCACACCCACGATCAACTGACCCATCGGTGTG
CTGATGTCGATTCCCTCGGTACACGACACCACGGTCTTGCCGTGATCTTCGCACCAGTCGCAGACCCTC
CTGAGCTGTCGGGCCTTCCGGCTGAGCCGATCTAGTTTCCAGACGACGATGATGTCGAACTCGTCGGCG
CGGTTGTTGAGCCAGTCGCCGAGCTGGGGAGTGTCGAACGGGTCCGTAGACCCGGACACGTCTATATCC
TCGGCCCAACCGACGACCGTGTGACCGTTGGCGTTGGCCCACTGCTCGATAACCTCACGCTGGCGCTCG
ATCGAGGTCGATTCCTCGGAGGCGACCGACAATCTGGCTCTTCCCAAGACTTTCATGCAGGTCATGGTA
CGTTAGATTTCGCACAACCGTAGATACGGGATCCCATATCTCTGATTGCGCGAAAACAGGCAGCCACTT
ACTGCAGGTGGGTGGCCAACACCGAAGGAGAATCGCACCGTGACCGCAGCACTCATCGACCTTGTAGAC
GGCCAGCTTTCGGAGATGGAGATCAGCGAACCAGGCTTCATGTACGCGGGCAGGCTGACCAATCTGGAG
ACCGGGGCATCGTGGCGGTTCGTCTACCGCAGCACGATCGTCAAAACCACCAGCCCTATCGGCAAGCGG
ATGCTGGCGTGGCTCGACGGCGCGGCCCCAATCGGGCAAGCAAAGATCGCCGCCGACCCCGACATGATG
TACAGCACCGCCGAGCTGACGCTCTGGGAGCCGTGAAAGGTGAACTGACAGCCGCACTCGACGCGCTCT
AATCTGAAGACTTCTTCCGGCAAGCCCCCCGCCAAGCGCGGGGGTTTTGTCGTCTGAAGGGCAGGAGGC
CAGCCTGATGGCCCCTCGTATGTGAGTCTCGCCACATCGAGGCCCAGATTCAGCAGATCAGACCCATAT
ACAAGTGAGAGACAAAGAGAGAGTCCCTGCGGGCGTAAGCGTGGTGTTCGTCACAAAGAGGCCCAGATT
CTGCAAATTACACGCATATACAAGTAGAGGGGTTCGAGAGAGCCCCGGAACGAACTTGCTGGCTTGAGA
GCGCGTACCGCGCCTCGTCAGCAGCCTGACGGTCGGGTGTCCTACTCCATCCTCCCCGACCGCCTGGTC
TTCCCCTGGTGTACCTGCGCCAGCGGTAGCCGTCCGAGTGGGTCGGACGGATAAGGCACGCCAGCCAGC
GTGCCGCCCTTTCTCTGTGGAGACCCCCGGTTGAGTCCATGAGGCCGACCGGGGGCTTTCCCGCCAGCA
CGCCAGAATCGCGTAATGCGTTCTGGCGCTTTTTATTTGACCTAAGCCATGAACGCACATGAAAGGAGG
GCACGATGCCCGCCACTACCGACCAGTGCCCTCAGGGGCACGAGATCCGGTCCACCGCCGACCGCGACA
AGCAGGGCTACTGCCGCCGATGCCGAGCTGACAACGCACGCAAGCGCCGGGTGGGCAAGGAGGCCGCGC
TGATCGTCGTACGCGCACTTGAGAAGGCCGGCGTTCAGTTCCAGAACGACGGCGTGCCCGCAGAGCCCG
CCGAGGTAGTACGCCAGCTCACCGACGCATACGCGGCTGGCTTGTTCGACCAGCAACCGCAGTAACCCG
GCTGCGAATCCCAGATGCCGTGCCCAGGAGGCAGGCAACCCGATTAAGGAGAACACGATGCACGACAAC
GACTTTGACCACGACCGCGCCCAGCGGCTGATACTCAACCTCAAGGTGGAGAACCGCAGGCTAAAAGCC
CGTTTGCGGGATGAGATCGGACGGCTCACAGACCAGGAGCTGCCGCCGTCCTGGCATAAGCGGTTCGCC
CAGCTACGCGTTGAGAACGCCAGGTACCGCATCGAGCGCAACGAAGCTCGCGCCGAGGTAGCACGGCTC
CGCGACGAGCTGGCAGCGGTCTCCAACGATGGCTAACCGCAACCCGGTCAAGCACCACGGCGTGCTCGG
CTCCCACGGACCTGGAGGAGGACGGTTGTTCGCCTCGGTTCCGATGGCGCTGGCTCGTGACACCGAGCT
ATCCGCGCCCGCCCGGTCGGTGGCCATGTTCGTCTGGTCGCACGACGAGAAGTACCAGCAGTCAGCCCG
TGACGTTGCGGACGCGTTCGGCATGAGCCGCAACACCGTGGCCAAGGCGCTCACGGATCTTCAGGAGCG
CGGCTGGTTCGTACGGGAGCGGATCACTGCTAACTCCGAGGTGTGGCACCTGCAGATGACCAACCGCCC
GTTCACCGCAGACGAGGTCTCGAAGCTGTCCAGGGTGGCTCAGGAATTGAGCCACCTACCAAGCACTGG
CTCAGAATCTGAGCCACCTACTGGCTCACCAAATGAGCCACCTACTGGCTCAGGAATTGAGCCACATAG
TAGTGAAAGCAGGAGTGCACCAGAAGTGCACGACTGCAGTAGTGCAAGGACTGGTCTAGACGAGACCGG
CCTTTGGAGCCGGTCTCTCGCGGATGCCGGAAGAACAAGAGCGGCTGAACCTGCATCCGCAGAAGACCC
GTTCGCGGGTCCGCAACCCGCTTGGGTCGTAGAGGCAGCCGCACGCGAGGAGCGTGCTTGGGAATCGCA
ACCCGCATCCGCAGGGGGCCTACCGCCCCTGCCGTGGGAGTGACAACCGAGCAAGCAACTGGAGCCCTG

GCCTACAGCCGGGGCTCTCGTCGTTAACGGGCTCGGCAACCGCCGCGCCTCAACCAAATGAGGAGGAGA
CACATGCAAGACCCAATCGCTGAGGCAGCCGCCAGGGCTGCCGAGTACGGCGCACCGCCGAAGACGTGG
CACGTCGACCCGGACGCTGCAGTCGAGTACGGCACCGGACGCAACCCGCTAGACGGGATGGTCGTGACT
CACTGGAGCCACGATGGCTGAGACGACGTTCGAGCAGTACCTCGAACGGTCTGAGCCCTACTTCGCAAG
CATCCGCGAGGCAGGAGACCTGCCGTGGTTCGAGGACCCCGACAGGAGCCGCACCGTTGCAGAGCGCCT
GGGACTGCCGTCCGACATCGACCCGCTGGAGCTGCGCCGCGCTCTTTGGCAGCGCAGGAGCTACACCGG
AGCGGCAGTGTGAGCGTTACTCCGGACTTCAAAGTCTCGGTGCCTCATTTCCTGCACGCCGCAGGAAAC
GGGCATCAGCTCATCGTGGACCGCTCGCTGGCGTTCCACGTCCGAGACGAGCGGTTCCGCCCCGTCCTG
GACTACATCGCCGAGCACGGCCTGGTCCTGTTCGAGCGCACGATCAAACAGATCGATGGGCTGTTCGGG
CGCGTCGAGGAGAGTTATTACGTGACGCCGGACATCCGCAACCTGCTTCTGACCAACATCGATGAGGAG
ACACCCGCATGACCGAGACGAACCACACAACTCTGCCAGAGGCTGTGAGCAACGCTGACGGCCAAACGG
ACACTCCCGAGGGTCCTGGTGCCGCAGAACAGAATCAGGGCCTTATAGAGGCGCGCGACCGATACCGCA
CCGAGCGTGACGCCGCGCGCGAGGAGCTGTCCACCGCTCATGCTCGAATCGAGCGGATGCAGCGAGCCG
AGGTCGAGCGGTTGGCAGCCGATGCCGGTCTGGCGATGGGTGGTGATCTGTTCATCAACGGCAACGCCG
TGGCTGACTACCTGACCGAGGACGGAGACGTGGACGCCGAGAAGGTGGCCGCTGATGTCGATGCAGTCC
TGGCCGAGCGGCCTGGTCTGCGAAAGAACGCACACGCGGTCGATCCGTCACAGGGGCGCGGTGGTCCAG
CCCCGAAAGCCTCTGAACCGACGTTCGCCGATCTCTTCAAGTCTTGAGCATGAGCCGTCCGTGACGGCC
TAGCTCACCAACGCCTCTGTGAGGCACCCCAAGGACTGACCTGTGGTCGGTCCTTTTTTGTTGCCCAAC
AACACGTTTCAAGGAGAAACACATGGCAATGCAGCACTCCACCACCGCCGATGCTTGGACTCCCGAGGA
TTTCGGCAAGGTTCTCAACAAGGCCATTCAGGCCCAGTCGACCGCGTTCCGCGCGGGCACCACATTCGG
CACTGACAAGGTCCGCGTTAGCTTCCCGCTGTGGAACGCCGACCCGTCCGCAGCCTGGCTCGATGAGCT
GGAGCTGATCGTTCCGACCGACGGTGCCACCGGCTCAGTCGTTTGCACCCCCTCCAAGGTTGGCGGCAT
CACGACGCTGTCGAACGAAATCCGCAACGACACCGACCCCGCGATTGCTGATCAGGCCGCCGCCGGCCT
GGCGAACGACATCGCGAAGAAGATCGACGCAGCGTTCTTGGCGAACACCACGGCCAAGGCGAACAACGG
CCTCCTGTCGTTGACGACTTCGGTCGTGGACACCGGCGCATCGATCGCGAACGAGGATCCGTTCATCGA
CGCGATCTTCGCCGCTGAGGCGGTCGGCGCGAACATCGACCGCTGGATCATGCACCCCGACACCGCCAA
GGTCCTGTCGAAGCTGAAGAAGGGCACCGGCTCGAATGAGCGGCTGTTGCAGCGCACCACCGACGGCGC
GTTCATCGCGGGCATCAAGGTTCTGACCGACCCCAAGGTCGACGTCAACACCAAGGCATGGGGTATCGA
CTCCACCCGCACCAAGATCGTGCTCCGTCAGGGCACCGAGATGCGGCTGTTCGACGTTCCGCGCCAGGA
CGCCATCGACGTTCGCGGTATCGCCCGCGTCGGCTTCGCGTTCCTGCATCCGGAGTCGGTTGTGCGTCT
GTACGACGCCGCCTGATTCATCCCTCGTAACCCTCACGGAGGGGCTGAGTTTCGTGCTCAGCCTCTCCT
CTGAGGCATCACCCCGAAGGAGGTGAACATGGCTAGAAAAGCTGGAATGTCGACCGGCGTCGAGGTCGC
TCGGATCTCGGTCAAGGTGTCCCCCGACACCAAGAACTTCCGACGCGAGCTGAAGAGGGATCTCGATCG
CATCGAGGAGTCGATGCGGGCCAAGATCGACGTAGAGCCCGACATGAAGGGGTTCCGGCAGCGGGTCAA
CGCTCAGACCAAGGGCATGCGGACCAGCGTCAAGGTCGACGCCGACGTTGACCGCAAGGGTTTCCTCGG
TCGGATCGCCAACAGCCTGAGCCAGATTCAGCCGCCGTCGTTCGGCAGCGGGATCAACCCGACCGGGTA
CGCCCTGATCGCGGCGGGCATCGCCGCGCTGGCACCACTCATCGCAGGCACGCTCGGAGCCGCCACAAC
GGCTCTGCTCGCGCTCCCTGGCCTGGCCGCTGCAGTCGCGACACCGATCGCCGCGCTGACGCTCGGTAT
CGACGGGCTCAAGAAGGCAGCCGAGACACTCAAGGATCCGTTCGAGGATCTGAAACAGTCGATGTCCTC
GGCGGTCAAGGATCAGTTCAGCCCGGTGTTCGAGCAACTCCGGTCGCTGTTCCCGACGCTGAAGGCTGC
ATTGCCTTCTGTCACAAAGGGTTTGGCCGACATGGCCAAATCGATTGCCGGTGTCATCACCTCCCCCGA
GGGGCTGGCCAAGATCGATACGACCATCCGCAACATCGGCACGGCGCTGACCACAGCCGCGCCAGGCGT
CGGCAAGTTCACCGAGGGCCTGATGGGCCTCGTGGAGTCCTTCACCGGCAAGCCCCTGCAGGGCGTAGC
CGACTGGTTCACCAAGACCGGAGATTCGTTCTCGGCGTGGGTCGAGAAGATGACCCGGCCTAGCTGGTT
CACCGGCAAGAGCCCGCTTGAGGTTGCGTTCGGCAACCTCGGCGACACGCTCAAGACCATCGCCGACAC

CCTCGGCGACGTAGGCCAGAAGGCTCTCGATTTCTTCTCCGACCCGGAGAAGGTGAAGAGCTTCAAGGA
CGAGCTGACGTTGCTGTCGGATGTCATCTCCGGAATCGCGACCGGCATCAACGGAATCGCCTCGGCGTA
CTCCAAGCTCCCGCTATCCGGCGAAGGTCTGAAAGGGCTTATGCCGATCCAGGCTCAGCTCGGAGTCAA
GATGTGGGACGGCCTGAAAGAAGGTGCGGCGAAAGCGTTCGCCGAAGTGTCCACGATGGCAGTCGGGTT
CGTCTCGAACATCGGCAGCACGTTCGCGAACATCGGCAACACGCTCTCGGGCATCTGGAACGGCGCGGT
CTCAGCCGCGCAGTCTGCGTGGTCCTCGATCGTCAGCACCGTGTCCGGTGCGGTCGGGAACGTGGTCTC
TAAGGTCACCGGCATGGTGGGTCAGATCGTCTCCACGCTCGCCAGCCTTGCAGCAGAGGGCATCAACGC
TGGCCGCAACCTGGTCCAGGGCCTGATCAACGGCATCAGCGGAATGATCGGCTCAGCGATCGCCAAGGC
GCGAGAGCTGGCCAGCGGCGTAGCCGATGCGGTGAAGGGATTCCTCGGGATCCACTCGCCGTCGAAGCT
GTTCACCGAGATCGGTGAGTACGTCGGTCAGGGCTTCGACAACGGGCTCCAGAGCCAGATAGCTCGGCT
CGGCAAGACCGCCAGGGCGATGGCAGAGACGGTGACCGACGAGTTCCACGGCGGGCTGAAGTTCGGAGC
TGACGGGTTCAGCACCGACAGCGACAACCCGATGCTGCAGGCCGCGGCCGGCCTCGCCAACGCACCGGT
CGACTTCGCGAAAGCGACTGGTAAGCAGTTCCTTTCGGACATCGGCATCTCGGGCGAAGGCGTGCTATC
CAGGGCTGTCTCCGAGGGCATCAACTACGTGTTCCAGATCGGCTCTGTCGATGAGGCGCTGTCCATCAA
GGACCGCACGGAGTCCAAGCAGGCACTGAGCGTCGTCGGACGCTGACGAATCGATGAGGACCCCCGGCT
GTCTGGCCGGGGGTCTTCTCGTCAACGACTAAGGAGGAGAACGGCATGCAAGACAGATGCGCCAGACCG
GGTTGCGTGCGGACAGTCAAGAACATCGACCGGCACCGCGCGTGCTCGGCAGTCTGCCAACACATCCTG
CTCCAGATCGAGAACGCAGACAGGCTCTGCAACGCGCTGGGCGTTGAGGGGCTGACAGGCCAGTACCGC
GACGTGATCACCGCGTTGTCTGACGCTTGGACCCGAGCCCAGGAGCTGGACTACCACCTCTACCAAGAG
GCGAAGAGCGTAGGTATGCGTGCGTCGGACTGGGAGAGCTTGAAGAGGGGCAAGGCTTCGCGCCCTCCG
AGCTGATGTCAGTTGCTGCTGATCGACTTGATGATTTCGATGATGATCCACACCGGAATCCATAGACCG
CAGGTCAAGATCGTCAGGATCAGATGCACACCCGCGTCAGACCCACCTCCGCGCACGTTGACGTTGTTG
TACACCGTGGCGGTTGCTGCGGGAGCTACAGCCTGACGATGCTCGGTCCACCTCACCCCGTCCCAGTAC
CGCAGGAGACCGGGTGAGTGCGGGTCTGGGTACCAGTTGGGTGCTGGTGGAGGTGCTGGCGCGGAGTAA
GTCGGTGGGACCGGCACGGTCGGGTAGTGCTCGGGTTGGCGGTAGCCGACTACCTCACCGCTGTACTGC
TCAGAGTCTTTTCCCCCAGGTGTGACCACGGCAGCAGTCTAGGACACACGTACCTGAATGAACTGACGT
TCAGACGGATGCGCGACACCCCCAGGGGGTTACCCCCTCCACCCCGGTCTCGACCGGTCGGTTA
